# Behavioral syndromes can reduce population density: behavior and demographic heterogeneity

**DOI:** 10.1101/073262

**Authors:** Bruce E. Kendall, Gordon A. Fox, Joseph P. Stover

## Abstract

Behavioral syndromes are widely recognized as important for ecology and evolution, but most predictions about ecological impacts are based on conceptual models and are therefore imprecise. Borrowing insights from the theory of demographic heterogeneity, we derived insights about the population-dynamic effects of behavioral syndromes. If some individuals are consistently more aggressive than others, not just in interspecific contests, but also in foraging, mating, and anti-predator behavior, then population dynamics could be affected by the resulting heterogeneity in demographic rates. We modeled a population with a boldness-aggressiveness syndrome (with the individual's trait constant through life), showing that the mortality cost of boldness causes aggressive individuals to die earlier, on average, than their non-aggressive siblings. The equilibrium frequency of the aggressive type is strongly affected by the mortality cost of boldness, but not directly by the reproductive benefit of aggressiveness. Introducing aggressive types into a homogeneous non-aggressive population increases the average per-capita mortality rate at equilibrium; under many conditions, this reduces the equilibrium density. One such condition is that the reproductive benefit of aggression is frequency dependent and the population has evolved to equalize the expected fitness of the two types. Finally, if the intensity of aggressiveness can evolve, then the population is likely to evolve to an evolutionarily stable trait value under biologically reasonable assumptions. This analysis shows how a formal model can predict both how a syndrome affects population dynamics and how the population processes constrain evolution of the trait; we suggest some concrete predictions.

## Introduction

Behavioral syndromes—within-individual and between-individual consistency in behaviors across time or ecological contexts (Sih et al., 2004)—have been uncovered in a variety of animals, including mammals, birds, fish, and a variety of invertebrates (Sih and Watters, 2005; Riechert and Hedrick, 1993; Gosling, 2001). In a species with a behavioral syndrome, an individual is classified as having a particular behavioral type (BT), often quantified along axes such as aggression, activity, sociability, or fearfulness. Behavioral syndromes indicate constraints on adaptive behavioral plasticity across ecological contexts (Sih et al., 2004): for example, an aggressive individual may be more effective at foraging, but more vulnerable to predation. Behavioral types are sometimes heritable (e.g., Dingemanse et al., 2002), such that BT frequency may change in response to selection.

Individual behavioral variation is now recognized as important for ecology and evolution (e.g., Dall et al., 2012); Sih et al. (2012) and Wolf and Weissing (2012) hypothesized a variety of ecological consequences of behavioral syndromes, including effects on population dynamics. Aggressive behavior might stabilize density-dependent equilibria, through a shift from scramble to interference competition (Sih et al., 2012); a similar effect may emerge if aggressive individuals cannibalize less aggressive individuals (Pruitt et al., 2008; Andersson et al., 2007). In contrast, increasing frequencies of aggressive individuals may destabilize an equilibrium if they have a faster “pace of life” (Réale et al., 2010), if temporal behavioral correlations introduce time lags in the population’s response to fluctuating environmental conditions (Sih et al., 2012), or if frequency-dependence leads to coupled oscillations of BT frequency and population abundance (Sinervo and Calsbeek, 2006). Finally, behavioral syndromes may affect the intensity of density dependence, with potential impacts on equilibrium density: interaction rates may increase if aggressive individuals are more active (Pintor et al., 2009) and hence encounter conspecifics more frequently (Sih et al., 2012), but they may decrease if the various behavioral types use different resources and habitats (Wolf and Weissing, 2012). Intriguing as these hypotheses are, few are based on formal population models (a notable exception is the study by Fogarty et al., 2011, showing how heterogeneity in a sociality syndrome could affect invasion speed), and thus we do not know the range of conditions over which they hold.

There is some empirical evidence suggesting that behavioral syndromes may modify demographic rates such as birth and mortality rates. For example, a review by Biro and Stamps (2008) found that aggressiveness and boldness were consistently associated with increased birth rate, and a meta-analysis by Smith and Blumstein (2008) found that aggression was positively associated with birth rate and negatively associated with survival. These studies involved small and idiosyncratic sets of species, so we cannot draw strong conclusions about the generality of the results. However, when such effects occur, behavioral syndromes will lead to among-individual variation in demographic rates, which has come to be called “demographic heterogeneity” in population dynamics (e.g., Fox et al., 2006).

Fortunately, there is a body of quantitative theory in population ecology showing that, depending on within-population correlation structure (Engen et al., 1998) and the underlying stochastic process, demographic heterogeneity can change the variance due to demographic stochasticity (Vindenes et al., 2008; Kendall and Fox, 2003). Furthermore, persistent survival heterogeneity (i.e., phenotypic variation that creates lifelong differences in instantaneous or annual mortality risk) in long-lived organisms can increase the population’s density-independent growth rate (Kendall et al., 2011), increase its equilibrium density (Stover et al., 2012), and reduce its extinction risk (Conner and White, 1999). In contrast, persistent birth rate heterogeneity alone has more limited effects on dynamics. The differential effects of heterogeneity in the two types of vital rates can be understood by recognizing that differences in survival accumulate multiplicatively with age: as a cohort ages, the more “frail” individuals tend to die off and the mean survival of the cohort increases. This “cohort selection” (Vaupel and Yashin, 1985) means that the expected survival (averaged across all age classes in the population) is greater than would be found in a population with the same baseline value but no heterogeneity in survival. Differences in birth rates, however, accumulate only additively, and cohort selection on birth rate will only occur if birth rate is correlated with annual survival rate.

Models of demographic heterogeneity lead us to expect that any behavioral syndrome that introduces persistent heterogeneity in survival will have impacts on the low-density population growth rate and on equilibrium abundance, in ways not addressed by Sih et al. (2012) or Wolf and Weissing (2012). Here, we develop a population model that incorporates a syndrome in which aggressive individuals are more successful at reproducing, but experience greater mortality (e.g., because of energetic costs of aggression, or because of greater exposure to predation). Using this model, we show four quite general results. First, the equilibrium BT fitness is directly controlled by the mortality cost of aggressiveness but is affected only indirectly by the reproductive benefit of aggression (via a parent-offspring correlation). Second, a population with a polymorphism for behavioral types will typically have a different density-dependent equilibrium than one made up entirely of the non-aggressive BT, and the polymorphic equilibrium is most often lower. Such effects on density could affect the species’ local extinction risk and influence on the ecological community. Third, if parents evolve to produce an offspring BT distribution that equalizes the expected fitness of both types (as has been found by Pruitt and Goodnight (2014) in a social spider), then the equilibrium population abundance will always be lower than that for a monomorphic, non-aggressive population. Finally, we show that selection on the strength of the aggression trait may lead to an evolutionarily stable value (stable ESS); while the resulting level of aggressiveness depends on details of model functions, the existence and stability of the ESS are nearly guaranteed if the mortality cost is an accelerating function of aggressiveness. By establishing quantitative links between population dynamics and behavioral syndromes we hope to open up new realms of empirical inquiry in both fields.

## Model description

For simplicity of exposition, we modeled a population with two phenotypes, using the subscripts *a* and *n* to identify aggressive and non-aggressive individuals, respectively. As has been documented in various species (e.g., three-spined stickleback; Huntingford, 1976), aggressive individuals can monopolize mates or good territories, and thus have a higher birth rate, *β*: all else being equal, *β_a_* > *β_n_*. The other component of the syndrome is that aggressive individuals are bolder in contexts that may increase their mortality risk; thus, we model non-aggressives as having death rate *µ* and aggressives as having death rate (1 + *γ*)*µ*, where *γ* > 0 is the additional risk born by the aggressive BT. Thus, aggressive individuals have a fitness advantage over non-aggressive individuals if

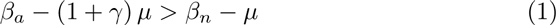

or

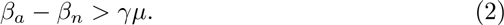

Table 1 provides a reference for all symbols used in the paper.

**Table 1:** Symbols used in this paper.

| Symbol | Definition |
| --- | --- |
| <u>Variables, parameters, and indices in model</u> |  |
| $N$ | Population abundance |
| $w_a$ | Frequency of aggressive BT<sup>1</sup> in population |
| $\pi$ | Frequency of aggressive BT among newborns |
| $\beta$ | Birth rate |
| $\mu$ | Death rate |
| $\gamma$ | Additional death rate of aggressive individuals |
| $\alpha$ | Intensity of aggressive trait |
| $a$ | Subscript to indicate the aggressive BT |
| $n$ | Subscript to indicate the non-aggressive BT |
| <u>Quantities and symbols used in model analysis</u> |  |
| $x^*$ | Asterisk: Superscript indicating that quantity $x$ is evaluated at demographic equilibrium ($dN/dt = 0$) |
| $q$ | Weighted difference in BT abundances: $q = N_n/(1 - \pi) - N_a/\pi$ |
| $\bar{x}$ | Overbar indicates the mean of quantity $x$ across the population |
| $N_0^*$ | Equilibrium abundance of monomorphic non-aggressive population |
| $\tilde{w}$ | Frequency of aggressive BT in population at demographic equilibrium when average birth rates equal average death rates |
| $\hat{w}$ | Frequency of aggressive BT in population at demographic equilibrium when differences between birth rates equal differences between death rates |
| <u>Quantities and symbols used in ESS analysis</u> |  |
| $R$ | Subscript indicating resident population |
| $I$ | Subscript indicating invader population |
| $B$ | Birth rate of invader in environment created by the resident population |
| $s_x(y)$ | Invasion exponent: per-capita growth rate of invader with phenotype $y$ in environment created by resident with phenotype $x$ |

<sup>1</sup>Behavioral type

If aggressive individuals always hold a fitness advantage, then, if there is an additive genetic component to the syndrome, we would expect the aggressive BT to become fixed. Thus, to model a population that maintains multiple BTs (as is often observed in natural populations), we must either invoke a genetic mechanism such as heterozygote advantage (which has not been demonstrated empirically), assume there is no heritable component to the syndrome (which contradicts empirical evidence), or assume that the fitness difference between the BTs varies with density and/or frequency. We adopt the latter assumption in our analysis. In particular, we adopt the plausible assumption that aggressive individuals lose reproductive fitness by interacting with one another (e.g., Pruitt and Riechert, 2009; Lichtenstein and Pruitt, 2015). Thus, defining the frequency of aggressive BTs in the population as

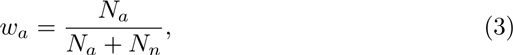

the birth rate of both BTs declines with *w_a_*:

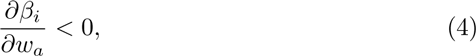

but that of the aggressive BT does so faster than that of the non-aggressive BT:

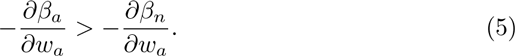

Finally, we assume that the birth rate of both BTs declines with density in the same way:

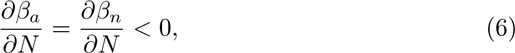

where *N* = *N_a_* + *N_b_* is the total population size. An example of birth rate functions displaying these qualitative features is illustrated in Figure 1.

**Figure 1:**
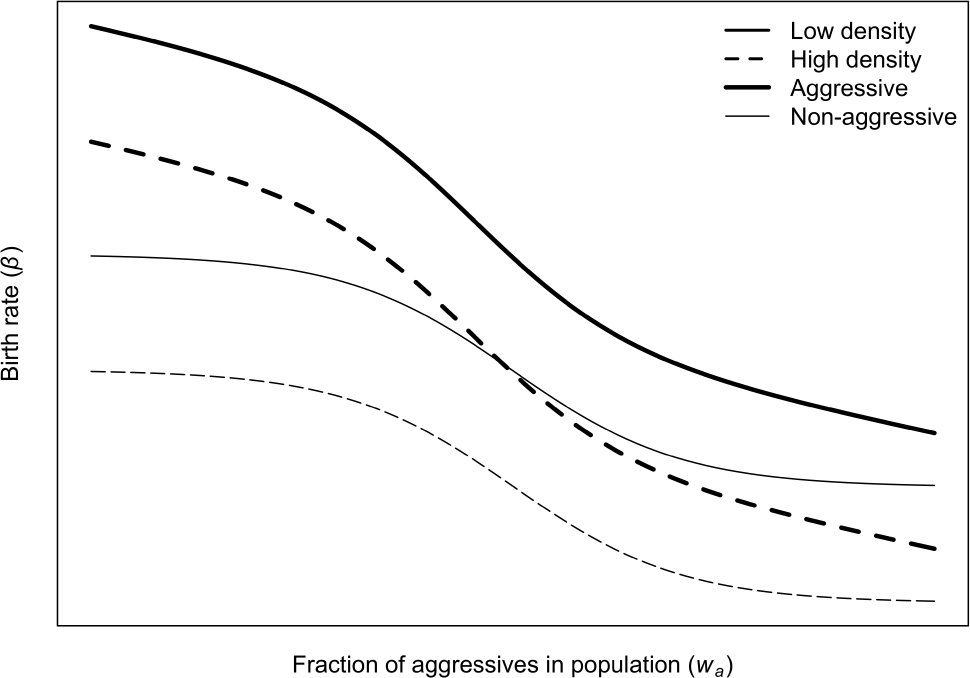
Example of density- and frequency-dependent birth rates that could arise from an aggression syndrome. Birth rates of both aggressive (heavy lines) and non-aggressive (thin lines) individuals decline with the frequency of aggressives in the population, but the relative advantage of aggression declines with increasing frequency of the aggressive BT. Birth rates of both behavioral types also decline with overall density (low density shown in solid lines, high density with dashed lines).

It is, of course, biologically plausible that frequency or density-dependence could instead (or in addition) occur in the death rate, or that the density-dependence is frequency dependent. We chose these particular assumptions to better draw upon the insights provided by the models in Stover et al. (2012). However, we do not expect that alternate assumptions about density- and frequency-dependence will qualitatively change our conclusions.

For the model to be explicitly about the boldness–aggressiveness behavioral syndrome, we must specify constraints and tradeoffs on the various functions and parameters. First, we let the birth rates depend on aggressiveness (*α*) as well as on the frequency of aggressives *w_a_* and density *N*, and assume that increasing the aggressiveness parameter increases the birth rate difference between the two BTs at a given frequency and density:

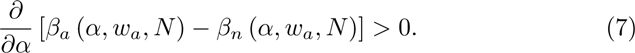

Furthermore, we expect that mortality is a function of *α*, so that increasing the aggressiveness parameter will also increase boldness, resulting in a greater mortality penalty:

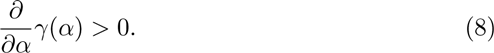

This tradeoff is needed to prevent runaway selection on the aggressiveness parameter.

We need one more component to build the population model: the frequency of each BT among the newborns. Behavioral syndromes have been demonstrated to be heritable (Dingemanse et al., 2002), but the underlying mechanisms have not been described. Therefore we assume simply that a certain fraction, *π_a_*, of an aggressive individual’s offspring are aggressive; 1 − *π_a_* of them are non-aggressive. Likewise, a fraction *π_n_* of a non-aggressive individual’s offspring are non-aggressive (*π_a_* and *π_n_* need not have the same value). The explicit functional forms of *π_a_* and *π_n_* depend on the details of the inheritance mechanisms, and even with simple two-sex genetic models (e.g., one locus and two alleles with dominance, or quantitative variation in an underlying latent trait) the functions will be quite complex (for example, if the behavior is controlled by an underlying continuous latent trait, *π_a_* and *π_n_* will depend on the frequency of aggressives in the population; Falconer, 1989). However, under a wide range of genetic mechanisms and mating systems, it is reasonable to assume that the rate at which a phenotype reproduces itself increases with the frequency of the phenotype; only when inheritance is near-perfect (e.g., parthenogenic with mutation or strong assortative mating) will the rates will be independent of phenotype frequency. Thus we assume

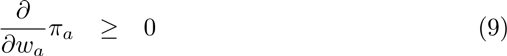

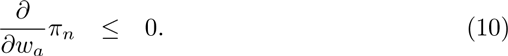

We can now write the population model:

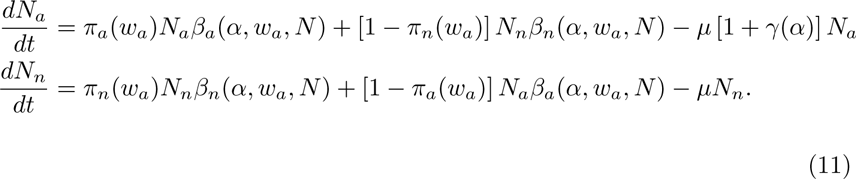

If the parent-offspring correlation is zero (as might arise if the behavioral phenotype is primarily controlled by environmental conditions that are uncorrelated between parents and offspring) then the fraction of newborns in each phenotype will be constant, and it will be more convenient to write

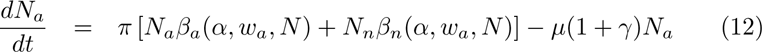

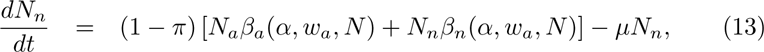

where *π* is the fraction of offspring with the aggressive BT.

These models differ structurally from the model of Stover et al. (2012) in three important ways: the flexible function for the fraction of newborns in each BT, which allows us to include both BT heritability and adaptive control of newborn BT frequency; birth rate functions that are both more flexible (the Stover model assumed linear density dependence in which the heterogeneity parameter affected both the slope and intercept) and allow for frequency dependence; and association of the “baseline” death rate with the non-aggressive BT rather than with the average phenotype. Nevertheless, many insights and analysis techniques can be carried over from Stover et al. (2012).

Note that some of inequalities (1–2) and (4–8) might be relaxed (become equalities) under special circumstances such as low density. However, it is reasonable to assume that they apply when the population is near its equilibrium density, which is where we conduct our analysis.

## Model analysis

There are a number of questions we want to answer about the model. First, for a given level of aggression, with associated boldness cost *γ*(*α*), what is the equilibrium frequency of the aggressive BT 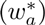? Second, is the associated equilibrium abundance (*N*^*^) greater or less than the equilibrium abundance that would be found in a population made up only of non-aggressive individuals 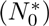? Third, is there an evolutionarily stable strategy (ESS) for the fraction of newborns that have the aggressive BT (*π*)? Finally, given a tradeoff between the benefits and costs of aggression, is there an ESS for *α*?

As written, equations (11) are too general to explicitly solve for *N*^*^ and 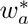. Even with the simplest form of the birth rate functions (linear dependence on density and frequency), the formulas for the equilibrium are too complex to provide much insight. However, we can get some qualitative (if sometimes vague) answers to the these questions by looking at the model from different perspectives. For example, rather than attempting to examine four-dimensional figures (birth rates as a function of total population size, the fraction of aggressives in the population, and the measure of aggressiveness), in Fig. 1 we examine a two-dimensional slice: for fixed *α*, we focus on how *w_a_*, the frequency of aggressives, might affect birth rates at two population densities. Much of the analysis below uses a similar heuristic approach.

### Equilibrium frequency of the aggressive BT

What is the equilibrium frequency of the aggressive BT in the population as a whole (which we denote 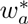)? First, let us look at the situation where the fraction of newborns that are aggressives (*π*) is constant. We can simplify the work by studying the variable *q* = *N_n_*/(1 − *π*) − *N_a_*/*π*, which is chosen because it allows us to eliminate the reproductive terms from eqs. (12) and (13) and to study the single quantity *q* rather than both *N_n_* and *N_a_*. The dynamics of *q* are given by:

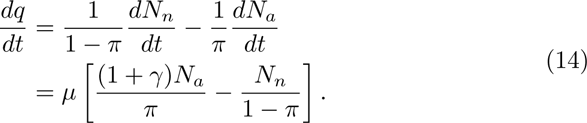

At equilibrium, 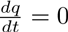; applying this to eq. (14) and rearranging gives

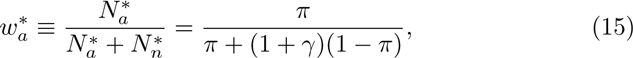

where the stars indicate that the model is being evaluated at equilibrium. Thus the aggressive BT’s equilibrium frequency depends only on its birth frequency and the mortality cost of boldness; increasing *γ* reduces the frequency of aggressives in the population relative to their frequency at birth (Fig. 2). This result is a consequence of cohort selection (Kendall et al., 2011): as a cohort of newborns ages, the aggressive BTs die faster, and so their frequency in the cohort declines. At equilibrium, the population growth rate is zero, and so the age and phenotype structures of the population match the life table of a cohort. Thus, the aggressive BT frequency in the population is found by averaging the frequency over all ages in a cohort, weighting by the fraction surviving to a given age, which will be less than the frequency at birth.

**Figure 2:**
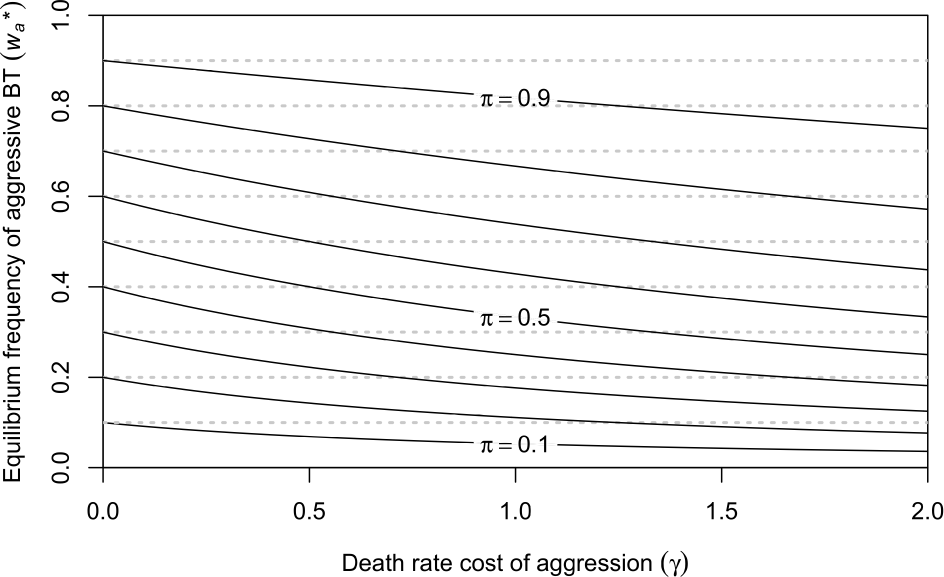
Equilibrium frequency of the aggressive BT in the population 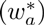 as a function of the mortality cost of aggression (*γ*). The curves are for different values of the aggressive BT frequency at birth (*π*), at equal intervals from 0.1 to 0.9.

Note that if the population is growing, then the age structure will tend to be biased towards younger individuals, relative to the equilibrium population. Younger cohorts have a higher frequency of aggressive BTs, since they have not been subject to so much cohort selection, and so a growing population will tend to have a higher aggressive BT frequency than will be found at equilibrium. By a similar argument, a population that is declining from above the equilibrium will have a lower aggressive BT frequency than the population will have at equilibrium.

When the phenotype is heritable, we can still analyze the model at the demographic equilibrium, where 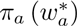 and 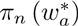 are constant. Here, we can write

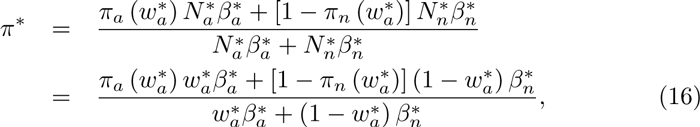

where 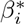 are the birth rates evaluated at 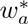 and *N*^*^. Inserting eq. (16) into eq. (15) and solving for 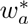 will give the equilibrium frequency. Unfortunately, for most inheritance functions this will not be analytically tractable, but it will still be true that the frequency of aggressive BTs will be lower in the population as a whole than among the newborns. In fact we can be more specific: at equilibrium, the average death rate in the population is the harmonic mean of the newborn death rates, as was shown by Stover et al. (2012).

Note that if a second gender carries the genes for the behavioral syndrome but does not express them, that gender will not be subject to cohort selection. Thus, the non-expressing gender will have a genotype frequency that matches that of newborns (and differs from that of the expressing gender). This substantially complicates the expression for the inheritance functions, but does not qualitatively change the fundamental result above.

### Aggression’s effect on the equilibrium population density

As a point of reference, we take the equilibrium density of a population made up of only non-aggressive individuals (which we call 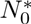), and we ask whether a population with both BTs has a population equilibrium that is larger or smaller than this reference. 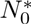 is defined as the density at which the non-aggressive birth rate matches its death rate: 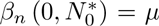 (in this section we are holding *α* constant so we suppress it for notational simplicity). If, near *w_a_* = 0, the aggressive BT’s birth rate is lower than its death rate, then the aggressive BT cannot invade the population, and so we focus on the situation where 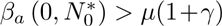. Increasing the aggressive BT frequency, *w_a_*, while holding 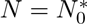 constant leads to declines in the birth rates of both BTs, but does not affect the death rates of either BT (because birth rates are density-dependent but death rates are not; Fig. 3). We can also define average birth and death rates:

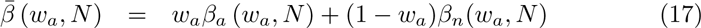

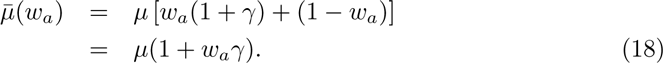

**Figure 3:**
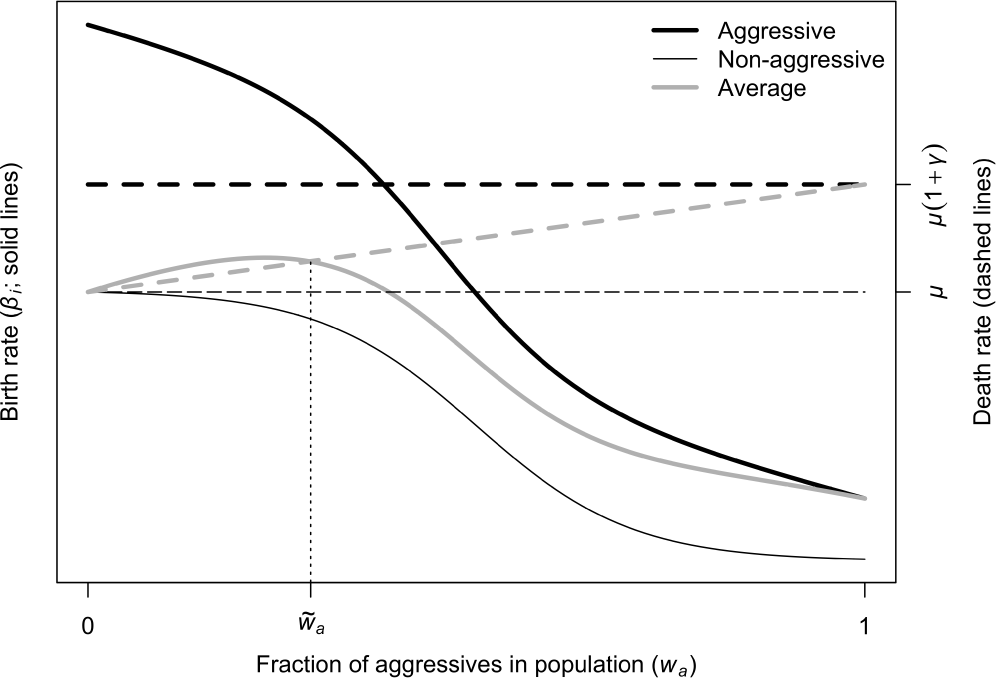
Birth and mortality rates of the aggressive BT (heavy lines), the non-aggressive BT (thin lines), and the population average (grey lines), as a function of *w_a_*, the frequency of aggressive BTs in the population. The overall population density is at the demographic equilibrium for a population made up of only non-aggressive BTs (at *w_a_* = 0, non-aggressive birth rate equals non-aggressive mortality rate). The vertical line indicates 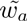, the positive frequency of aggressives that would reach demographic equilibrium at the same density (average birth rate equals average mortality rate). If 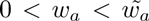 then the average birth rate exceeds the average mortality rate and the population would grow until it reaches demographic equilibrium at a higher density (lowering the birth rate curves; see Fig. 1). If 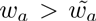 then the average birth rate is less than the average mortality rate and the population would decline until it reaches demographic equilibrium at a lower density.

While the death rates of each BT are constant, the average death rate increases linearly with *w_a_* because the aggressive type has a greater death rate. The average birth rate may show more complex patterns, because birth rates can also be density-dependent. In general, given constraints (5) and (6), the average birth rate will be maximized for a positive value of *w_a_* and its derivative with respect to *w_a_* will be greatest at *w_a_* = 0, as shown in Fig. 3. At 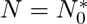, the average birth and death rates are equal at *w_a_* = 0. If the aggressive BT enters the population at low frequency, (so *w_a_* > 0), then, given the model assumptions there are three general cases we might see.

**Case 1**. First, the average birth rate might be greater than the average death rate for all values of *w_a_*. This would require that the average birth rate increase quite rapidly with *w_a_*. In particular, inspection of Fig. 3 reveals that this case requires that the birth rate increase faster than the death rate when the aggressive BT is rare and that the aggressive BT’s birth rate exceeds its death rate even when *w_a_* = 1. This can be shown to require:

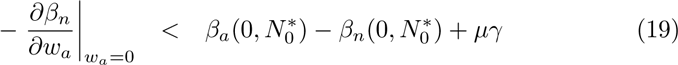

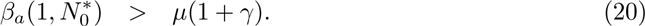

**Case 2.** The average birth rate might be less than the average death rate for all values of *w_a_*. This would require that that the non-aggressive birth rate decline sufficiently rapidly with *w_a_* when the aggressive BT is rare. In particular:

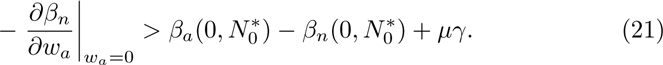

**Case 3.** The average birth rate might be larger than the average death rate for small *w_a_* and be less than the average death rate for large *w_a_* as is illustrated in Fig. 3. The average birth and death rates are equal at an intermediate value of *w_a_* which we call 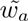.

In all three cases, if the population is found at BT frequency *w_a_* and total abundance 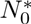, then abundance will increase if 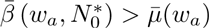 and decrease if 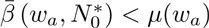 (of course, *w_a_* will then change dynamically as well). Now, we know that at equilibrium, the equilibrium BT frequency is 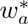, defined by eq. (15). We also know that at equilibrium, where *N* = *N*^*^, the average birth and death rates must be equal. Therefore, it is only possible for 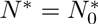 under the conditions of case 3 and when 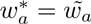. If 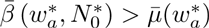, then 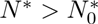; likewise, if 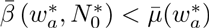, then 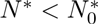.

Thus, the introduction of the aggressive behavioral syndrome into a naive population will increase the equilibrium abundance only if the aggressive BT has a very strong fitness advantage (taking into account the boldness cost) even at high frequencies (case 1) or if the equilibrium frequency of the aggressive BT is relatively low (case 3). Under case 2 or most circumstances of case 3 the syndrome will reduce the equilibrium abundance.

### Evolution of the birth frequency of the aggressive BT

For a given set of demographic parameters and inheritance functions, 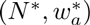 is the demographic equilibrium of the population model. However, it will not, in general, eliminate the fitness differences between the two BTs. To see this, suppose that case 3 applies and that 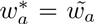, so that the demographic equilibrium is at 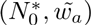, as shown in Fig. 3. At this point, the average birth rate equals the average death rate. However, the birth and death rates are not equal for either of the BTs. In particular, the non-aggressive BT must have negative net fitness, because its net fitness when 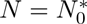 is zero only when *w_a_* = 0 and declines with increasing aggressive BT frequency. To achieve zero net fitness at the population level, the aggressive BT must have positive fitness. At the demographic equilibrium, the population growth rate is zero, and the reproductive value at birth for each BT is its birth rate divided by its death rate. At the equilibrium in Fig. 3, these are unequal. Just as Fisher (1930) showed with regard to the evolution of primary sex ratios, there will be selection on parents to increase the frequency of the type with the higher reproductive value (the aggressive BT) among their offspring.

If the inheritance function is purely genetic, with a fixed genetic architecture, then there is no way to respond to selection at this particular demographic equilibrium. However, if there are environmental influences (expressed directly or via epigenetic mechanisms) on an individual’s BT, then parents may be able to increase the frequency of the aggressive BT among their offspring, effectively changing the inheritance function (for example, androgen levels in egg yolk can influence offspring behavior, although this has not yet been explicitly linked to a behavioral syndrome; Ruuskanen and Laaksonen, 2010). Increasing the frequency of aggressives among newborns *π* will, in turn, increase the demographic equilibrium 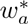 to a value greater than 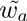. As shown in the previous section, this will lead to an equilibrium density that is less than 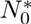. As long as the birth rates of the two BTs respond in the same way to density, at this new equilibrium, then, under the conditions of case 3, there will be a 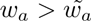, which we call 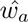, that satisfies eq. (1) – that is, the differences between the birth rates match the differences between the death rates. At the associated demographic equilibrium then not only do births match deaths for the population as a whole but also for each of the BTs (Fig. 4). Further increases in *w_a_* lead to a fitness advantage for the non-aggressive BT, so 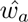 is the evolutionarily stable BT frequency. An analogous argument applies in case 2.

**Figure 4:**
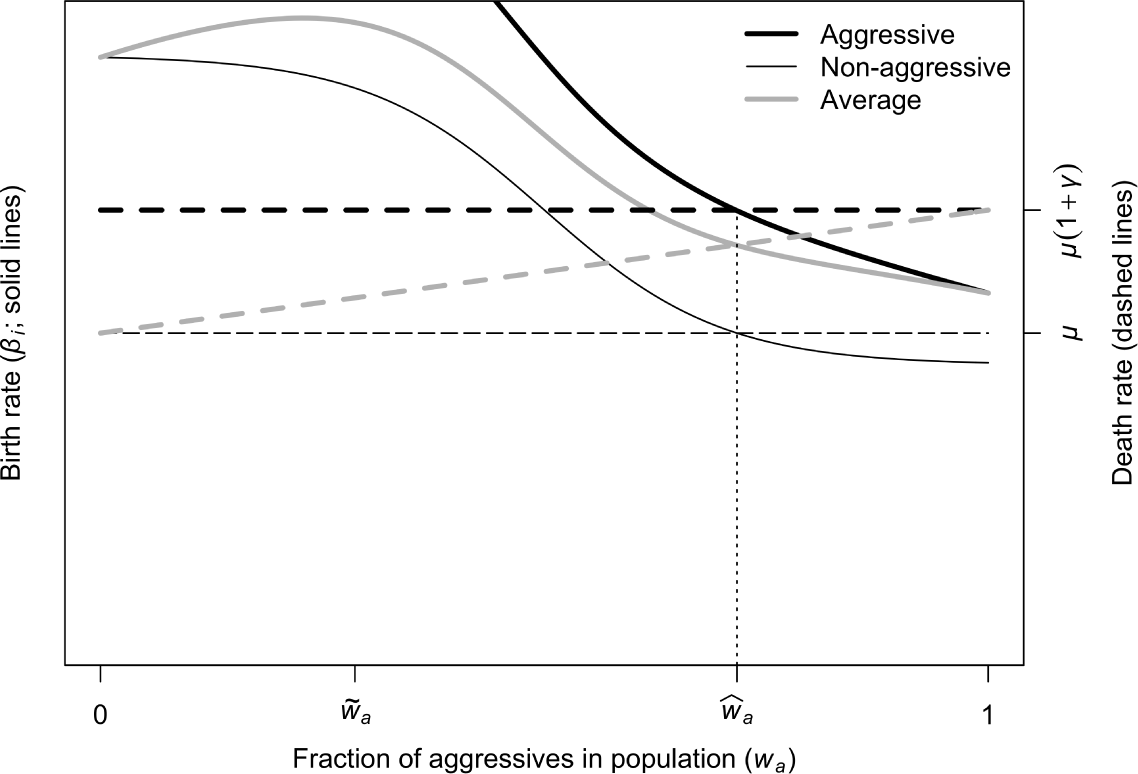
Birth and mortality rates of the aggressive BT (heavy lines), the non-aggressive BT (thin lines), and the population average (grey lines), as a function of *w_a_*, the frequency of aggressive BTs in the population. The overall density is at the demographic equilibrium for 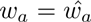, the frequency of aggressives at which the two BTs have equal relative fitness (the difference in fecundities equals the difference in mortalities). Because 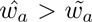, this demographic equilibrium is at a density lower than that of the non-aggressive-only equilibrium, allowing the frequency-dependent fecundities to be elevated.

In contrast, in case 1, where 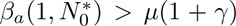, there is no frequency at which the two BTs have equal fitness at a demographic equilibrium (mathematically, 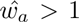), so the evolutionary stable BT frequency is 1 (fixation of the aggressive BT). Only in this last situation (at which we would no longer recognize a syndrome, as there is no behavioral variation) would evolution to a stable BT frequency result in a demographic equilibrium density that is larger than the density of a non-aggressive population 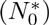.

### ESS for aggressiveness

In addition to the BT frequency, the strength of the aggressiveness trait (*α*) might itself be subject to selection. Incorporating an explicit genetic model for *α* would add a great deal of complexity to the model, so we instead take an adaptive dynamics approach, and look for an evolutionarily singular strategy (ESS) for aggressiveness (Geritz et al., 1998). In particular, we focus on the conditions allowing a resident population (with a given aggressiveness parameter, *α_R_*) that is at demographic equilibrium can be invaded by a population with a different aggressiveness parameter (*α_I_*). We start by stating the results, and then give the mathematical derivation.

To our existing model, we need to add one new quantity: the birth rate of an aggressive individual with trait *α_I_* in the presence of a given abundance and BT frequency of a resident population with trait *α_R_*. We call this 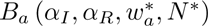; note that it will not be the same as the resident birth rate, 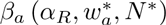. We also assume that the resident evolves to a birth rate frequency that equalizes the fitnesses of the two BTs, so that 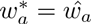.

The evolutionarily singular strategy, *α*^*^, is the value of aggressiveness that satisfies the condition

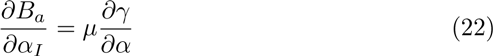

when evaluated at *α_I_* = *α_R_* = *α*. In other words, from the invader’s perspective, the birth rate benefits of increased aggression are exactly matched by the death rate costs when the invader and resident traits are identical.

The ESS is “ESS-stable,” meaning that a resident population that is at the ESS cannot be invaded, (Geritz et al., 1998) if

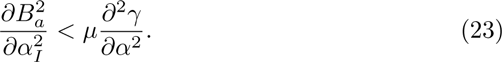

In particular, under the biologically reasonable assumption of diminishing returns to reproduction from increased aggression (making the left hand side negative), this condition will always be met if the the mortality cost is linear or accelerating in the aggression trait.

However, ESS-stability does not guarantee that the ESS can be reached through successive mutations of a resident population that is not at the ESS. This requires an additional property, called “convergence stability” (Geritz et al., 1998), and the ESS is called a convergence stable strategy (CSS; Diekmann 2004). Unfortunately, the formal conditions for the ESS to be convergence stable in this model require additional information, such as the relative sensitivity of the invader and resident birth rates to changes in the resident BT frequency and the inheritance mechanisms that determine the BT frequency in the invader population. While the condition can easily be calculated if functional forms are assumed, the general expression is sufficiently complex as to be non-informative. However, it seems clear that, if = 0 is not an ESS (that is, a mutant with slight aggressiveness can invade a population with none), then the existence of one or more ESSs at positive values of will ensure that at least one of them is convergence-stable.

### Mathematical derivations

The analysis of an ESS focuses on the the invasion exponent *s_x_*(*y*), which is the low-density per-capita growth rate of an invader with trait value *y* in the environment created by a resident population at equilibrium and having trait value *x* (the notation here follows Geritz et al. 1998 and Diekmann 2004). In the present context, *x* = *α_R_* and *y* = *α_I_*. The analysis proceeds by looking at derivatives of *s* evaluated at *y* = *x*. In particular, the condition for *x* to be an ESS is

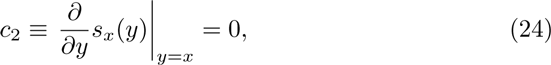

and the condition for ESS-stability is

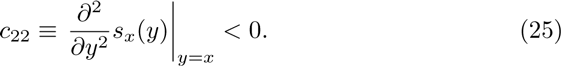

To calculate the invasion exponent we write out the dynamics of the invader population. Since the invader is rare, we assume that only the resident population *N*^*^ impacts the invader’s reproduction and that individuals of both populations are unaffected by the invader’s aggressiveness or BT frequency:

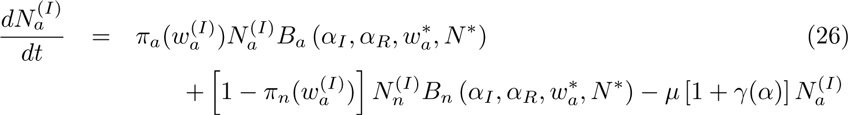

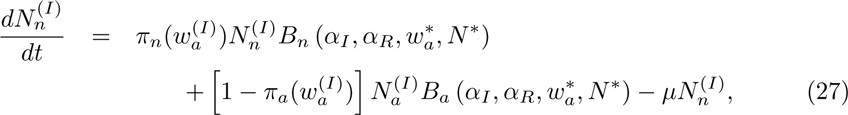

where 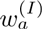 is the aggressive BT frequency among invaders (which might affect the BT frequency at birth). Adding these together and dividing by 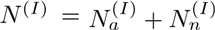 gives

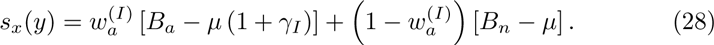

It is quite reasonable to assume that the invader non-aggressive BT is identical to that of the resident, so that *B_n_* = *β_n_*. Furthermore, if we assume the resident is at the fitness equalizing frequency, such that *β_n_* = *µ*, then the second term is zero, leaving

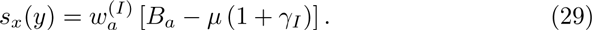

The first derivative is

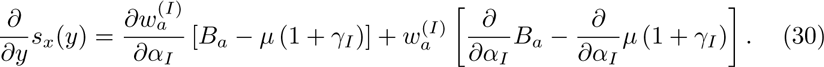

When *α_I_* = *α_R_*, it is quite reasonable to assume that *B_a_* = *β_a_* and *γ_I_* = *γ_R_*. Again, if we assume the resident is at the fitness equalizing frequency then *β_a_* = *µ* (1 + *γ_R_*) and the first term is zero. As long as 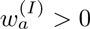, this leads directly to the ESS condition in eq. (22).

The second derivative is

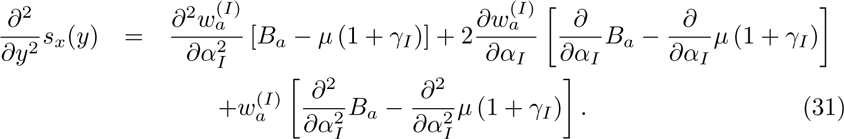

As above, when *α_I_* = *α_R_*, then the first term is zero. Furthermore, if the resident is at an ESS, then the second term is also zero (the quantity in brackets is just *c*_2_). As long as 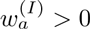, this leads directly to the ESS-stability condition in eq. (23).

## Discussion

We have developed a model that links behavioral and population processes by noting that the fitness associated with a particular behavior translates into demography—birth and death rates—at the population level. Applied to behavioral syndromes, this allows us to draw on existing theoretical frameworks (demographic heterogeneity and adaptive dynamics) to make a number of predictions about ecological and evolutionary outcomes, including the factors that control the frequency of the syndrome in the population, the effect of the syndrome on equilibrium abundance, and how selection should drive the evolution of both the distribution of the syndrome among offspring and the overall intensity of the behavioral trait. We applied the model to the boldness–aggression syndrome, but the general approach should apply to any syndrome with identifiable fitness consequences.

In the boldness–aggression syndrome, phenotypes differ in both their birth rates and their death rates. To a scientist not steeped in demographic theory, it may come as a surprise that these two effects are not commensurate. For example, we showed that when the population growth rate is zero, the frequency of the two behavioral types depends only on the frequency of the types among newborns and the relative death rates of the types—not on the differences in birth rates. We hasten to add that if there are genetically, epigenetically (e.g., Francis et al., 1999; Weaver et al., 2004) or environmentally induced correlations between the behavioral types of parents and their offspring, then the differences in birth rate will have an indirect effect via their effect on the distribution of newborn types. This occurs because mortality heterogeneity results in cohort selection, in which the composition of a cohort changes as the cohort ages (Vaupel and Yashin, 1985), whereas birth rate heterogeneity does not. One way of developing some intuition about this is to think about lifetime reproductive success (LRS) in a simple life history in which a phenotype’s birth (*β*) and death (*µ*) rates are both age-independent. Here, for an individual with phenotype *i*, the expected LRS is simply *β_i_L_i_*, where *L_i_* is the phenotype’s expected longevity. In a population with heterogeneous birth rates, the mean LRS can be found by using the mean of the phenotype-specific birth rates, and is unaffected by the amount of heterogeneity. Likewise, in a population with heterogeneous death rates, the mean LRS can be found by using the mean of the phenotypic-specific longevities. However, an individual’s expected longevity is an inverse function of its mortality rate: *L_i_* = 1/*µ_i_*. This nonlinear relationship means that the average longevity in the population will not be the same as the longevity of an individual with an “average” mortality rate; Jensen’s inequality (Zens and Peart, 2003) tells us that heterogeneity in *µ* will cause the population mean longevity to be larger than the longevity with the mean death rate, with the discrepancy increasing as the magnitude of the heterogeneity gets larger. This translates directly into effects on mean LRS. This fundamental distinction between birth and mortality heterogeneity persists even with age- and environment-dependent vital rates, as long as there is some degree of within-individual correlation through time (as in behavioral syndromes—and many other behavioral traits as well).

We found that the boldness–aggression syndrome often leads to a reduction in equilibrium abundance relative to a population made up of only non-aggressive types. This occurs because of the intersection of two factors. First, as the frequency of the aggressive BT increases, the mean death rate in the population also increases, reflecting the higher risk associated with boldness. Second, increasing the aggressive BT frequency increases the mean birth rate (for a fixed abundance) when the aggressive BT is rare (because the aggressives have higher birth rates), but the frequency-dependent depression of the birth rate drives down the mean birth rate when the aggressive type is more common. Once the frequency is high enough that the mean birth rate (at the non-aggressive equilibrium density) is below the mean death rate, then an equilibrium can only be reached if the density-dependence in the birth rate is relaxed by settling to a lower abundance. Note that, while our model incorporates density- and frequency-dependence in the birth rate only, qualitatively similar results would obtain if one or both dependences were in mortality. This affect on equilibrium density means that the behavioral syndrome may increase the population’s local extinction risk, with implications for conservation and metapopulation dynamics. It may also reduce the species’ trophic and facilitative impacts on other members of the ecological community.

Even at the demographic equilibrium, the two BTs will not necessarily have equal fitness. When they do not, there will be selection to increase the proportion of the more fit BT. If there is a response to this selection (whether through plasticity or evolutionary change), the BT frequency will move towards a value where both BTs have the same fitness. We have shown that this frequency equilibrium will always be at a value that results in a reduced equilibrium abundance, relative to a purely non-aggressive population. In density-dependent populations, selection maximizes the equilibrium abundance (often thought of as the carrying capacity, *K*), if fitness is not frequency dependent (Charlesworth, 1980). But because of the frequency-dependence of fitness of each BT in our model, behavioral evolution reduces abundance and thus increases the risk of population-level extinction due to stochastic fluctuations or exclusion by a competitor that can persist at lower resource densities (Webb, 2003).

How does aggressiveness (*α*) evolve? Our results show that there may be an evolutionarily stable value, but predicting where that will occur requires an understanding of the fitness of an invader with one aggressiveness level in a population of residents with a different aggressiveness. It may be reasonable that the outcome of the interaction between two aggressive individuals with different levels of aggressiveness only depends on the difference between the two *α*’s. Thus, the left hand side of eq. (22)—the derivative of the invader birth rate with respect to the invader aggressiveness, evaluated where the invader and resident have the same aggressiveness—will be independent of the resident aggressiveness level. However, changing the resident aggressiveness level will also, in general, change the resident’s equilibrium density and BT frequency; by analogy to the analysis comparing polymorphic and monomorphic populations, we might expect that increasing aggressiveness will decrease *N*^*^ and increase *w*^*^. We cannot say much in general about how these will impact the invader birth rate, especially as the predicted changes are likely to have opposite effects. If the effects balance out, then the left hand side of eq. (22) will be a constant, and an ESS will only exist if the right hand side is non-constant—that is, the death rate cost is a nonlinear function of aggressiveness. Turning to the stability condition, it is reasonable to assume that the invader birth rate has diminishing returns to aggressiveness, making the right hand side of inequality (23) negative. Thus, if the mortality cost is an accelerating function of aggressiveness, then both the existence of the ESS and its stability will be guaranteed.

We have modeled the behavioral syndrome as a dichotomous trait, in large part for ease of analysis and exposition, but a trait such as aggressiveness may take on a continuous range of values. In simple models of demographic heterogeneity (without density-dependence, frequency-dependence, or inheritance), a continuous distribution of death rates has been shown to have virtually identical effects on population dynamics as a dichotomous trait, the key value being the variance of death rates (Kendall et al., 2011). In models with density dependence, a key value for two-type models is the harmonic mean death rate among individuals in a newborn cohort (Stover et al., 2012); one might expect this to generalize to continuous trait distributions, but this has not been tested. Frequency-dependence in continuous traits has to be handled with care; the simplest approach is to assume a strict hierarchy, so that the fitness of an individual with aggressiveness trait *α_i_* only depends on the frequency of individuals with traits greater than *α_i_*.

Some qualitative predictions that derive from our model are described in Table 2. Nevertheless, applying this theoretical framework to particular species will require explicit functional forms for the density- and frequency-dependence in vital rates, as well as behavioral effects on these functions. These can be estimated empirically in focal populations, but further generalization will require a more mechanistic understanding of the underlying physiological bases of behavior (which generates the behavioral correlations) and the factors that link behaviors to fitness. The latter obviously include the risks of injury or death from inter- and intra-specific interactions, but could also include processes such as energetics (e.g., aggressive individuals might have a higher metabolic rate that forces them to be more risk-tolerant during foraging). A predictive theory also needs a mechanistic basis for the newborn phenotype distribution—e.g., what are the roles of genetics, parental effects, plasticity? Where genetics are known, we would need to develop explicit inheritance models using population genetics (Charlesworth, 1980) or quantitative genetics (Barfield et al., 2011); the latter would probably be most appropriate when the behavioral trait is continuously distributed. These are all areas where empirical research needs to guide model development. Such models could be used to understand the causes and consequences of phenomena such as the reduced fitness of heterotypic matings across a boldness syndrome in guppies (Ariyomo and Watt, 2013).

**Table 2:** Qualitative predictions about populations exhibiting a boldness-aggression tradeoff that derive from the theory developed in this paper.

| Prediction | Rationale |
| --- | --- |
| Compared with a population at a BT frequency that equalizes fitness between the types, an artificial population made up solely of non-aggressive individuals will, under identical environmental conditions, grow to a larger equilibrium abundance. | Corollary of the result that introducing an aggressive BT to a non-aggressive population will lower equilibrium abundance at the fitness-equalizing BT frequency. |
| The aggressive BT in populations growing in low-predation environments will exhibit stronger aggressive tendencies than the aggressive BT in high-predation environments.<sup>1</sup> | Reduced boldness cost at a given aggressiveness level will allow evolution to higher aggressiveness. Assumes local evolution of aggressiveness trait. |
| Populations in low-predation environments will have a higher aggressive BT frequency and may, depending on the strength of the boldness cost and the contribution of predation to overall mortality, have a lower equilibrium abundance than high-predation populations.<sup>1</sup> | Low predation reduces $\mu$, reducing mortality difference between BTs; this moves the fitness-equalizing frequency to a greater fraction of aggressives, where the birth-rate difference is also lower. For a given density, this reduces mean birth rate; if baseline mortality is not much affected by predation then reduced birth rate may exceed reduced mean death rate, requiring reduced density to attain demographic equilibrium. Assumes local adaption to fitness-equalizing BT frequency. |
| Populations in which aggression is under sexual selection will have an increased BT frequency and reduced equilibrium abundance relative to otherwise identical populations without sexual selection | Sexual selection increases difference in birth rates without affecting difference in death rate; thus fitness-equalizing BT frequency will be a greater fraction of aggressives. Effect on equilibrium density follows directly from this. Assumes no differences in aggressiveness trait. |

Quantitative predictions will require a model that is tailored to the empirical system being studied. Deviations from the model’s qualitative assumptions about density- and frequency-dependence will require new analysis to confirm that the predictions apply.
<sup>1</sup> Assumes that the boldness cost is a consequence of heightened vulnerability to predation.

The population model used here is very simple; in particular, it does not have age- or size-structure, and assumes that environmental conditions are constant. These factors can generate time-lags in feedback loops (because of the time to reach maturity) and fluctuating selection (e.g., fluctuations in predator populations, or in populations of alternate prey, which might lead to fluctuations in the relative cost of boldness as well as in overall mortality rates), respectively, and so may prevent the population from settling down to an ecological or evolutionary equilibrium. However, except in long-lived species, these are likely to be second-order effects that primarily affect quantitative rather than qualitative predictions. Of greater import, the model does not account for differences between sexes. Some species exhibit sexual dimorphism in behavioral traits (e.g., Pruitt et al., 2011; Han et al., 2015), and while individuals of the sex that does not express the syndrome do not experience the direct fitness effects, they may still influence the phenotypes of their offspring via genes or parental effects. If the frequency of aggressive BTs is below the level where both phenotypes have equal fitness and there is a genetic contribution to behavior, then individuals of the non-expressing sex could increase their fitness by preferentially mating with aggressive individuals. This would put the syndrome under sexual selection, and would further increase the birth rate advantage of aggression without necessarily increasing the boldness cost (Logue et al., 2009).

A recent essay on “data-free papers” in the behavioral syndromes literature (DiRienzo and Montiglio, 2015) suggests that such papers (encompassing syntheses of older theories as well as novel conceptual frameworks, terminologies, or statistical approaches) are contributing relatively little to our understanding of the subject. Notably absent from their critique are formal models; this is perhaps due to their relative paucity (we have found only one model linking behavioral syndromes to population dynamics; Fogarty et al. 2011). However, DiRienzo and Montiglio (2015) suggest (correctly, in our view) that formal models are an important avenue (along with empirical study) to effectively study the speculative links suggested by verbal conceptual frameworks. The work here represents such a contribution. In particular, our findings that the equilibrium frequency of the aggressive behavioral type depends strongly on the mortality cost of boldness and that the equilibrium population abundance is negatively related to the frequency of the aggressive BT could only have been derived from a model that translates the fitness consequences to the individual into birth and death rates, and included dynamic feedbacks via density-dependence and frequency-dependence. Indeed, these feedbacks help shape the fitness landscape in which the behavioral syndrome evolves in such a way that understanding the optimal values of the intensity of the syndrome and the BT frequencies requires explicit modeling of the species’ population ecology. Making quantitative predictions about specific systems will require tailoring the model to those systems; such species-specific models can then be used, for example, to predict and explain patterns observed in common garden experiments with animals drawn from different selective environments. We hope that this paper will stimulate studies that integrate empirical observation and formal modeling.

## Acknowledgements

This work was supported by the National Science Foundation (grant numbers DEB-1120865, DEB-1120330). We thank Jonathan Pruitt for comments on the manuscript.

